# Sustained pupil responses are modulated by predictability of auditory sequences

**DOI:** 10.1101/2020.11.10.376699

**Authors:** Alice Milne, Sijia Zhao, Christina Tampakaki, Gabriela Bury, Maria Chait

**Affiliations:** Ear Institute, University College London, London WC1X 8EE, UK; Now at: Department of Experimental Psychology, University of Oxford, Oxford OX1 3PH, UK

**Author notes:** Corresponding Author: Alice Milne.

## Abstract

The brain is highly sensitive to auditory regularities and exploits the predictable order of sounds in many situations, from parsing complex auditory scenes, to the acquisition of language. To understand the impact of stimulus predictability on perception, it is important to determine how the detection of predictable structure influences processing and attention. Here we use pupillometry to gain insight into the effect of sensory regularity on arousal. Pupillometry is a commonly used measure of salience and processing effort, with more perceptually salient or perceptually demanding stimuli consistently associated with larger pupil diameters.

In two experiments we tracked human listeners’ pupil dynamics while they listened to sequences of 50ms tone pips of different frequencies. The order of the tone pips was either random, contained deterministic (fully predictable) regularities (experiment 1, n = 18, 11 female) or had a probabilistic regularity structure (experiment 2, n = 20, 17 female). The sequences were rapid, preventing conscious tracking of sequence structure thus allowing us to focus on the automatic extraction of different types of regularities. We hypothesized that if regularity facilitates processing by reducing processing demands, a smaller pupil diameter would be seen in response to regular relative to random patterns. Conversely, if regularity is associated with heightened arousal and attention (i.e. engages processing resources) the opposite pattern would be expected. In both experiments we observed a smaller sustained (tonic) pupil diameter for regular compared with random sequences, consistent with the former hypothesis and confirming that predictability facilitates sequence processing.

**Significance statement:** The brain is highly sensitive to auditory regularities. To appreciate the impact that detecting predictability has on perception, we need to better understand how a predictable structure influences processing and attention. We recorded listeners’ pupil responses to sequences of tones that followed either a predictable or unpredictable pattern, as the pupil can be used to implicitly tap into these different cognitive processes. We found that the pupil showed a smaller sustained diameter to predictable sequences, indicating that predictability eased processing rather than boosted attention. The findings suggest that the pupil response can be used to study the automatic extraction of regularities, and that the effects are most consistent with predictability helping the listener to efficiently process upcoming sounds.

## Introduction

The sensory environment is laden with regularities. The brain readily exploits this predictable information, using it to drive perceptual experiences (de Lange et al., 2018), guide attention (Zhao et al., 2013) and influence decision-making (Soltani and Izquierdo, 2019). In the domain of hearing, our ability to use these statistics plays many important roles, from auditory scene analysis (Bendixen, 2014; Heilbron and Chait, 2018) to discovering regularities in the speech signal (Erickson and Thiessen, 2015).

Accumulating work demonstrates that listeners automatically detect predictable structure in unfolding sound sequences. In a seminal demonstration, Saffran et al (1996) showed that infants are able to segment a continuous stream of syllables based only on the statistical relationships (frequency of co-occurrence) between adjacent elements. This paradigm has since been expanded to a variety of statistical structures and behavioral tasks to reveal robust “statistical learning” across the life span (Conway, 2020). Sensitivity to statistical regularities is also exhibited in the brains of naïve listeners during passive exposure to sound patterns (Barascud et al., 2016; Southwell et al., 2017) and in other species (Milne et al., 2018; Wilson et al., 2017).

A key question pertains to understanding how the detection of predictable structure influences processing and attention. The link between regularity and attention has been contentious. On the one hand it is argued that regularity automatically biases attention (Mackintosh, 1975; Itti and Baldi, 2009; Feldman and Friston, 2010; Zhao et al., 2013; Alamia and Zénon, 2016), consistent with the premise that regular structure in the environment carries important information about behaviorally relevant elements within our surroundings, and should therefore receive perceptual priority and attentional resources. On the other hand, a large body of work demonstrates that the brain exhibits reduced responses to regular, predictable stimuli (de Lange et al., 2018; Richter et al., 2018), interpreted as reflecting the fact that the detection of regular structure facilitates the conservation of processing and computational resources. Indeed, it has been shown that regular patterns are easier to process (Rohenkohl et al., 2012) and also, critically, easier to ignore (Andreou et al., 2011; Southwell et al., 2017; Makov and Zion Golumbic, 2020) which has been taken as evidence that regularity does not draw on attentional resources.

Here we use pupillometry to tap into these different cognitive processes. Pupil diameter is a commonly used measure of bottom-up driven salience and processing effort. Non-luminance-mediated pupil dynamics are controlled by a balance between norepinephrine (NE), reflecting the activation of the arousal system (for reviews see Joshi et al., 2016; Larsen and Waters, 2018)and acetylcholine (ACh), hypothesized to correlate with the processing load experienced by the individual (Sarter et al., 2006). By studying pupil responses to structured vs. random auditory patterns we sought to determine how sustained pupil diameter, and by proxy the listeners’ arousal and processing load, change as a function of regularity.

If regularity facilitates processing, a smaller pupil diameter would be predicted in response to regular relative to random patterns. Conversely, if the emergence of regularity is associated with an increased demand on attention, we expect the opposite pattern - a larger pupil diameter associated with more predictable stimuli, reflecting increased salience-evoked arousal and a consequent draw on processing resources.

We studied two types of predictable acoustic structure: in Experiment 1 we used deterministic (i.e. fully predictable; Figure 1) sequences, as described in Barascud et al (2016), to study the pupil response to regular, relative to randomly-ordered, tone pip sequences. These sequences were generated anew on every trial, tapping into processes that rapidly detect, and exploit, the predictable structure. In Experiment 2 we used a more complex probabilistic structure similar to the classic Saffran paradigm (Figure 2). These sequences did not follow a deterministic order, instead the transitional probabilities between tones allowed the stream to be segmented into triplets. Listeners were pre-exposed to such sequences, and pupil responses were measured subsequently to quantify responses to the pre-acquired statistical pattern.

**Figure 1.**
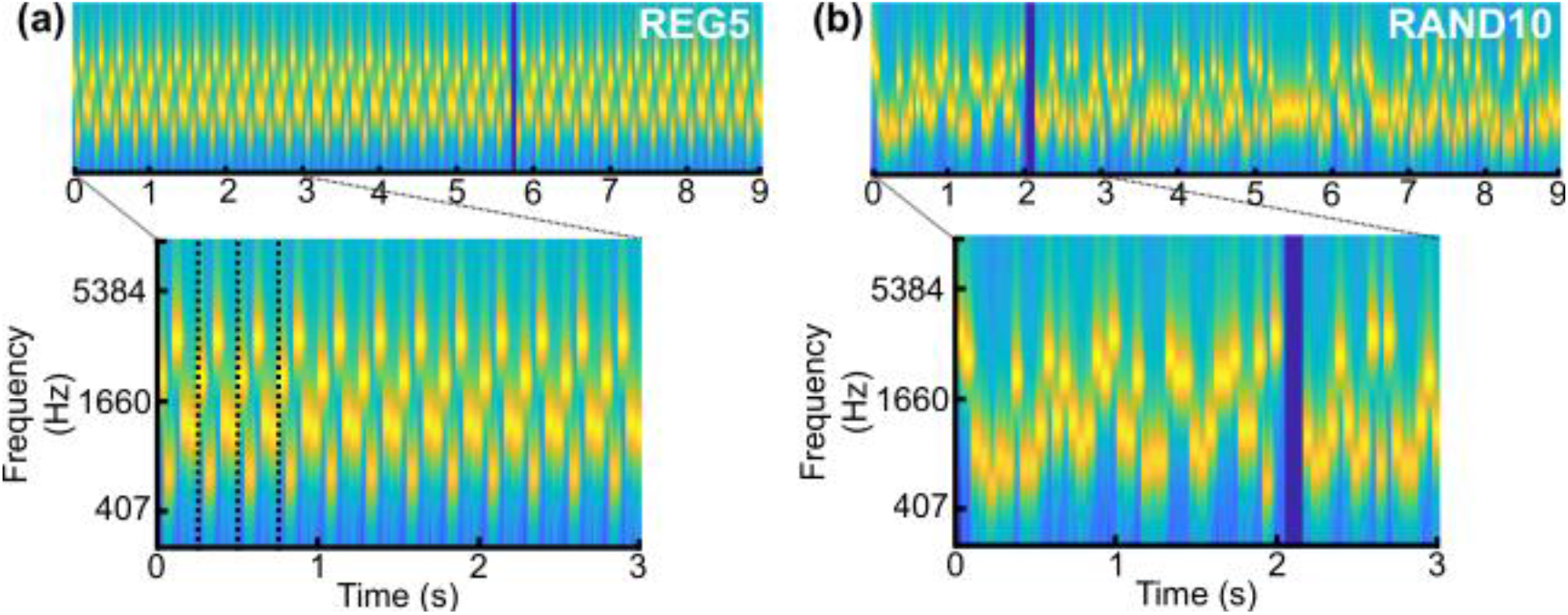
Stimuli used in experiment 1. Stimuli were sequences of contiguous tone pips (50ms) with frequencies drawn from a pool of 20 fixed values. The tone pips were arranged according to frequency patterns, generated anew for each subject and on each trial. REG sequences were generated by randomly selecting 5 (REG5), 10 (REG10) or 15 (REG15) frequencies from the pool and iterating that sequence to create a regular repeating pattern, (**a**) example of a spectrogram for REG5, dotted lines indicate the first 3 cycles. RAND sequences were generated by randomly sampling 5 (RAND5), 10 (RAND10) or 15 (RAND15) frequencies with replacement. (**b**), example of a spectrogram for RAND10. A subset of trials were target trials containing a gap generated by the removal of 2 (REG) or 3 tones (RAND), indicated by the dark blue band in the spectrogram.

**Figure 2.**
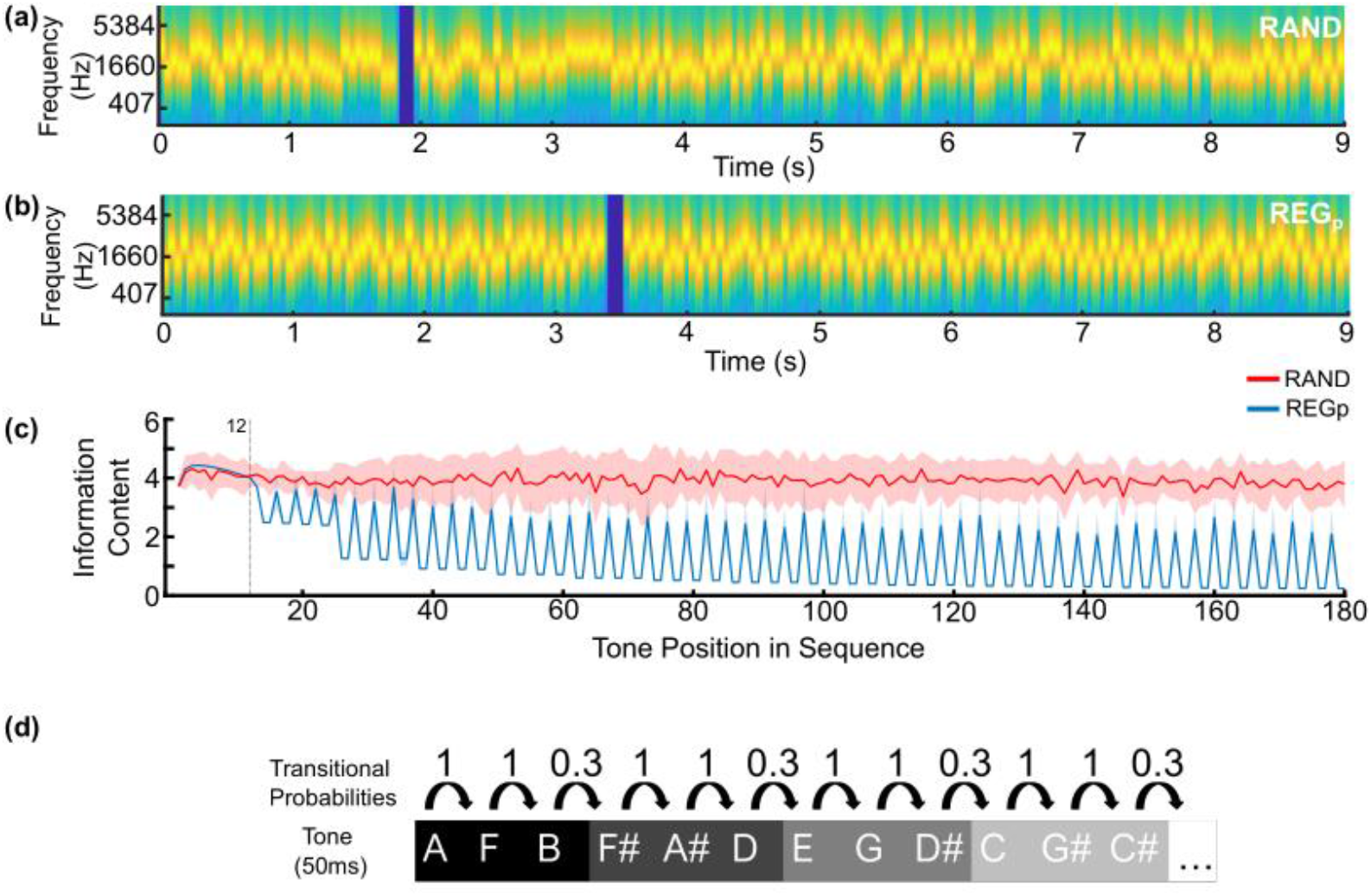
Stimuli used in experiment 2. Stimuli were sequences of concatenated tone pips (50ms) with frequencies consisting of 12 different tones that corresponded to the musical notes shown in (**d**). (**a**) spectrogram of RAND sequences where the tones do not follow a predictable pattern. A subset of trials were target trials containing a gap generated by the removal of 3 tones and is indicated by a dark blue band in the spectrogram of **a** and **b**. (**b**), spectrogram of the “regular” condition generated from a probabilistic pattern shown in (**d**, top row); in the regular condition (REGp), tones were arranged into four three-item tone ‘words’, the four words are shown in different shades of gray. The tones within a word always occurred together giving them a transitional probability (TP) of 1. Each word could transition to any of the other words, giving tones at word boundaries a TP of ∼0.3. As a result, these sequences do not have a regular structure in the same way as experiment 1, compare with Figure 1a. **(c)** Ideal observer model response to RAND (red) and REGp (blue) signals shows the information content of each tone pip (averaged over 24 different sequences). The output of the model shows the information content (IC) for each tone as the negative log probability (−log P) of a tone pip and therefore the higher the IC value the more unexpected the tone. The model indicates that while IC remains consistently high for unpredictable sequences (RAND, red), for REGp (blue) it begins to drop after 12 tones (i.e. after each ‘word’ has repeated). Evidence for the probabilistic structure then continues to accumulate throughout the sequences as indicated by the gradual separation between the REGp and RAND ICs. Shading indicates ± 1 SEM. (**d**, bottom row). The random sequences presented the same tones as the regular sequences but in a random order.

## Materials and Methods

Results from two experiments are reported. We continuously tracked pupil diameter while participants listened to 9-second-long sequences of contiguous tone pips, that either contained a predictable structure or did not. To control participants’ attention, and to make sure it was broadly focused on the auditory stimuli, an incidental, easy gap detection task was used; listeners were required to monitor the stream of tones and indicate when they noticed a silent ‘gap’ within the sequence. The gaps, generated by the removal of several consecutive tones, were placed at a random position in ∼25% (experiment 1) and 20% (experiment 2) of the sequences. Participants were kept naïve to the presence of an underlying pattern to enable the study of implicit sequence learning. This study was not pre-registered.

### Stimuli and Procedure

Participants sat with their head fixed on a chinrest in front of a monitor (24-inch BENQ XL2420T with a resolution of 1920×1080 pixels and a refresh rate of 60 Hz), in a dimly lit and acoustically shielded room (IAC triple-walled sound-attenuating booth). Sounds were delivered diotically to the participants’ ears with Sennheiser HD558 headphones (Sennheiser, Germany) via a Roland DUO-CAPTURE EX USB Audio Interface (Roland Ltd, UK), at a comfortable listening level that was adjusted by the participant during the practice phase. Stimulus presentation and response recording were controlled with Psychtoolbox (Psychophysics Toolbox Version 3; Brainard, 1997) on MATLAB (The MathWorks, Inc.).

#### Experiment 1

Stimuli were 9-second-long tone sequences (Fig. 1a and b) of contiguous 50ms tone pips (ramped on and off with a 5 ms raised cosine ramp; 180 tone pips per sequence). Tone frequencies were selected from a pool of 20 logarithmically spaced values between 222-2000Hz. Sequences were generated as previously described in Southwell et al. (2017). A unique sequence was presented on each trial. Sequences were defined by two parameters: regularity (whether they consisted of a regularly repeating or random pattern) and alphabet size – the number of frequencies comprising the pattern (5, 10 or 15). In regular (REG) sequences, a subset of frequencies (‘alphabet size’) were randomly drawn from the full pool and arranged in repeating cycles. Paired random (RAND) sequences were generated for the same frequency subset by randomly arranging the tones. Therefore, REG and RAND conditions were matched for the occurrence of each frequency. Overall six conditions were used (RAND/REG x 3 alphabet sizes; REG5, RAND5, REG10, RAND10 and REG15, RAND15).

Approximately 25% of the stimuli contained a single silent gap anywhere between 1 and 8 s after sequence onset. This was created by removing two tones from REG sequences (100ms gap) and three tones from RAND sequences (150ms) to equate task difficulty (Zhao et al., 2019b).

The experiment consisted of seven blocks (∼ 8 mins each) and a practice block. There were 24 trials per block (4 trials per condition) for a total of 168 trials (28 trials per condition). Inter-trial intervals were jittered between 2500-3000ms.Stimuli were presented in a random order, such that on each trial the specific condition was unpredictable.

Throughout the block a black cross was presented at the center of the screen against a gray background. Participants were instructed to fixate on the cross while monitoring the sequence of tones for gaps, and to respond by button press as quickly as possible when a ‘gap’ was noticed in the tone stream. At the end of each trial, visual feedback indicated whether gaps were detected correctly. Further feedback was given at the end of each block, indicating the total number of correct responses, false alarms, and average response time. The practice block contained six gap trials (3 REG, 3 RAND) to ensure participants understood the task. In the main blocks only 25% of the trials contained gaps. The experimental session lasted approximately 2 hrs. A break of at least 3 minutes was imposed between blocks to reduce the effects of fatigue.

Previous work with MEG (Barascud et al, 2016) and EEG (Southwell et al., 2017; Southwell and Chait, 2018) demonstrated that brain responses in naïve passive listeners rapidly differentiate RAND from REG signals, with responses to REG diverging from RAND within 2 regularity cycles. We expected pupil responses to also follow this pattern and show a change in pupil size once the structure has been acquired. Further, we expected the change in pupil size to occur later for larger alphabet sizes, as more information is required in order to identify a longer pattern.

#### Experiment 2

Experiment 2 investigated sequences that contain a probabilistic rather than deterministic structure. Sequences were based on the pure tone version of the segmentation paradigm introduced by Saffran and colleagues (Saffran et al., 1999), with the key modification, that instead of the 333ms long tones in Saffran et al (1999), we used 50ms tones.

To generate the underlying probabilistic structure twelve different tones were arranged into four tone ‘words’ made from the following musical notes, AFB, F#A#D, EGD#, CG#C# (Fig. 2d), these corresponded to frequencies: A = 440 Hz; A# = 466.16 Hz; B = 493.88 Hz; C = 523.25 Hz; C# = 554.37 Hz; D = 587.33 Hz; D# = 622.25 Hz; E = 659.25 Hz; F = 698.46 Hz; F# = 739.99 Hz; G = 783.99 Hz; G# = 830.31 Hz. As in Saffran et al. (1999) the same tone ‘words’ were used for each subject. Sequences were generated anew for each trial by randomly ordering the tone words, with the constraint that the same word did not occur twice in a row, thus tone words always transitioned to a different tone word. This created a probabilistic structure where the transitional probability (TP; the probability that tone “a” will be followed by tone “b” calculated as the; frequency of a *to b*/frequency of *a*) between tones within a word was 1, and the TP at word boundaries was 0.33. RAND sequences were generated in the same way as for experiment 1 but using the 12 frequencies listed above.

To formally demonstrate how this probabilistic structure emerged over the course of a sequence we used a PPM (prediction by partial matching) statistical learning model. The model, Information Dynamics of Music (IDYOM; Pearce et al., 2010), uses unsupervised statistical learning to acquire the transitional probabilities of tone pips within each sequence. The output of the model shows the information content (IC) for each tone as the negative log probability (−log P) of a tone pip and therefore the higher the IC value the more unexpected the tone. The model output (Fig. 2c) demonstrates that, following presentation of the first 12 tones (each of the four tone ‘words’), the two types of sequence, regular (REGp, blue) and random (RAND, red) rapidly diverge. While the random sequences remain unpredictable, the tones in REGp gradually become more predictable as the model learns the sequence structure. In contrast to deterministic regularities (see model in Barascud et al., 2016), these probabilistic sequences have a much more gradual change in information content. As a result we would expect that for this, more complex, regularity the information tracked by listeners will be less precise, and thus likely to have more variability across individuals. For this reason, we introduced a familiarization phase to ensure listeners had ample opportunity to become sensitive to the structure. This familiarization phase consisted of only REGp sequences. Participants were then tested on REGp and RAND sequences while recording the pupil response. Following pupillometry measurements, a further behavioral test was administered to more explicitly probe if the subjects had become sensitive to the regularities. Therefore experiment 2 consisted of the following three phases:

1. **Familiarization:** The familiarization phase gave listeners ample opportunity to acquire the probabilistic structure. In this phase, trials consisted of 27-seconds-long REGp sequences (540 individual tones in total) such that each ‘tone word’ was encountered 45 times within each sequence. A gap detection task was used to ensure participants attended to the sequence. Due to the length of the trials, each sequence contained two gaps. The gaps were generated by removing 6 tones, creating a 300 ms gap. The gap was intentionally longer in the familiarization phase to make the task easy and reduce the effects of fatigue for the next phase. Overall, the familiarization stage lasted ∼7.5 mins consisting of 15 trials. Participants were instructed to respond (key press) when they heard a gap. After each trial participants received visual feedback on the number of correct responses and false alarms. No pupil data were collected in this phase.
2. **Pupillometry:** Following a minimum three minute break, participants completed the pupillometry phase. All trials contained a 9-second-long tone sequence (180 tones in total, 60 tone words). 20% of trials (“target trials”; REGp and RAND with equal proportion) contained a single gap that occurred between 1 s and 8 s post-onset. In all conditions, the gap was 150ms long (removal of three tones). This phase consisted of two blocks of 30 trials. This provided a total of 24 trials per condition.
3. **Behavioral probe:** This phase tested how much knowledge listeners had gained about the structure of the sequence. Pupil responses were not recorded. We conducted two separate probes designed to test familiarity and sensitivity to sequence structure. In the Familiarity probe, participants were presented with sixty 3 second trials (REGp vs. RAND; 50% of each condition). They were instructed to listen carefully to the sounds and decide if the sequence felt “Familiar” based on the initial exposure phase. They were told to use a ‘gut’ feeling if they were unsure. In the Structure probe, participants were instructed to listen and identify if the sequence contained any sort of structure, or, appeared to be random. The two probes were completed by the “main” group (those participants who completed the Familiarization and Pupillometry stages), and by a “control” group that was recruited to only complete the behavioral probes. The purpose of this control group was to establish the degree to which the structure could be extracted without prior exposure. As these participants had no prior exposure to the REGp and RAND stimuli in the familiarity probe they were told to use a ‘gut’ feeling to identify familiar sequences.

### Participants

#### Sample size

We aimed for a sample size of approximately 20, based on previous data from a related pupillometry study (Zhao et al., 2019a) where robust pupil response effects were observed using as few as 10 participants.

#### Experiment 1

22 paid participants were recruited, four were excluded providing a final sample size of 18 participants (11 females, mean age 25.2, range 19-35). In both experiments, exclusion occurred either during data collection e.g. due to difficulty tracking the eye or excessive blinking or tiredness (eye closure), or due to a high blink rate that was identified in pre-processing, before separating trials by condition.

#### Experiment 2

For the main group, 24 paid participants were recruited, four were excluded providing a final sample size of 20 participants (17 females, mean age 21.2, range 19-28). The control group consisted of 20 paid participants (10 females, mean age 22.3, range 18-30).

All participants declared that they had no known otological or neurological conditions. Experimental procedures were approved by the research ethics committee of University College London and written informed consent was obtained from each participant.

### Pupil diameter measurement

An infrared eye-tracking camera (Eyelink 1000 Desktop Mount, SR Research Ltd.) was positioned at a horizontal distance of 65 cm away from the participant. The standard five-point calibration procedure for the Eyelink system was conducted prior to each experimental block and participants were instructed to avoid head movement after calibration. During the experiment, the eye-tracker continuously tracked gaze position and recorded pupil diameter, focusing binocularly at a sampling rate of 1000 Hz. Participants were instructed to blink naturally during the experiment and encouraged to rest their eyes briefly during inter-trial intervals. Where participants blinked excessively during the practice block, additional instructions to reduce blinking were provided. Prior to each trial, the eye-tracker automatically checked that the participants’ eyes were open and fixated appropriately; trials would not start unless this was confirmed.

### Statistical Analysis

Statistical analysis was conducted in SPSS (IBM SPSS Statistics, version 27) and Matlab (Mathworks, 2017a).

#### Behavioral Data

##### Gap detection task

For experiment 1, sensitivity scores (d’) were computed using the hit and false alarm rate (z(hits) -z(false alarms). A keypress was classified as a hit if it occurred less than 1.5 s following a target gap. Where hit rates or false alarms were at ceiling (values of 1 and 0, respectively; resulting in an undefined d’) a standard correction was applied whereby 1/2t (where t is the number of trials) was added or subtracted. For four out of six of the conditions d’ was not normally distributed, therefore Wilcoxon signed rank tests were used to compare REG vs RAND performance. We first averaged d’ across alphabet sizes to test the main effect of regularity (REG vs RAND). As there was a main effect of regularity, we then conducted three pairwise comparisons (Wilcoxon signed rank) to test if the effect was present for all alphabet sizes. We were not interested in the effect of alphabet size independent of regularity therefore did not test this as a main effect. P-values were adjusted for multiple comparisons using the Holm–Bonferroni method. For experiment 2, no false alarms were made, therefore only Hit rate (HR) was computed and analyzed. Due to normality-violating ceiling effects Wilcoxon signed-rank tests were again used to compared REGp vs RAND performance

Reaction times (RT) were recorded from each ‘hit’. For experiment 1 these were analyzed with a repeated measures (RM) ANOVA with factors of regularity (REG vs RAND) and alphabet size (5,10,15). For experiment 2, a paired-samples t-test was used to contrast RAND and REGp. Reaction times met the assumptions for parametric tests and alpha was a priori set to *p* < .05. An additional exploratory RM ANOVA was conducted to compare reaction times that occurred early (< 4.5s) or late (> 4.5s) in the trial. Regularity (REG vs RAND) and time (Early vs Late) were entered as factors. No post hoc tests were run for this analysis.

##### Behavioral probe (experiment 2 only)

For the two probe tasks, sensitivity scores (d’) were computed as described in the previous section. To test if d’ scores were higher in the main group relative to the control group, who were naïve to the sequences, an independent samples t-test compared group (‘main’ vs ‘control’) for each probe task. Spearman’s correlations were used to test if performance (d’) for the two probes (familiarity vs structure) was correlated across the two tasks. For each probe, exploratory analysis also correlated d’ against pupil diameter for each time point in the trial (down-sampled to 20hz), using Spearman correlation. We present the correlation coefficient at each time point and indicate time points where p < 0.05, family-wise error (FWE) uncorrected.

### Pupillometry data analysis

Trials containing a gap and trials where the participant made a false alarm were excluded from the analysis. Most participants made infrequent false alarms in experiment 1 (see figure 3-1), and only 3 subjects made more than one false alarm per condition. Between 17 and 21 trials were analyzed per participant per condition ([20-21] for REG5, REG10, REG15; [19-21] for RAND5; [17-21] for RAND10). There were no false alarms in experiment 2.

**Figure 3.**
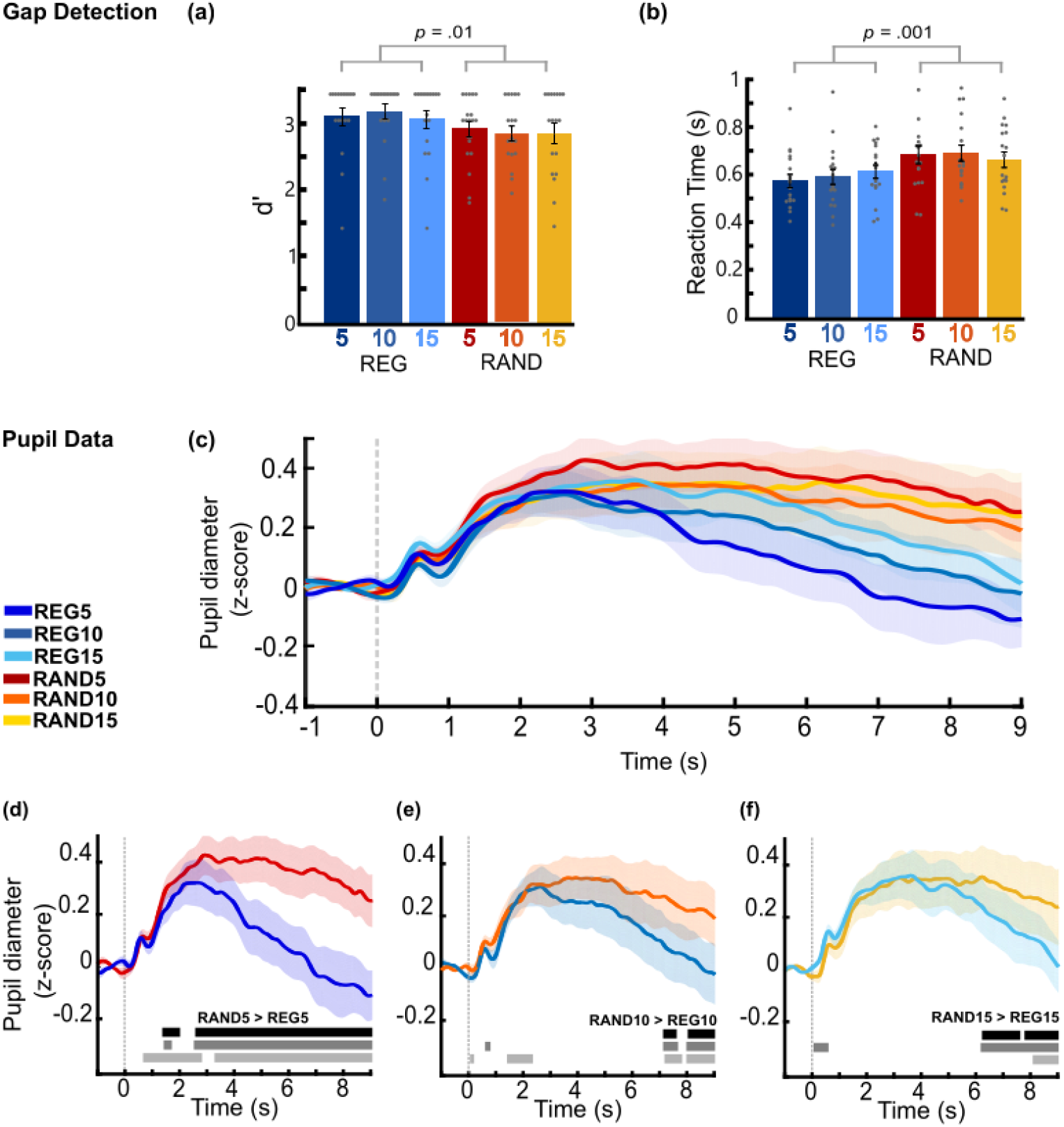
Experiment 1 – Regularity modulated pupil size and gap detection performance. The gap detection task showed worse performance for RAND compared to REG sequences. Performance measures were: (**a**) dprime (d’) and (**b**) reaction time (RT) for each of the six conditions, see Figure 3-1 for hit rate and false positive data. Sensitivity to the gap was significantly higher, and RT shorter for REG relative to RAND sequences. Circles represent individual data points. Error bar shows ± 1 SEM. Plots (**c-f**) show averaged normalized pupil diameter over time, baseline corrected (−1 – 0s pre-onset). The shaded area shows ±1 SEM. The horizontal bars show time intervals during which significant differences (bootstrap statistics) were observed. The black bar shows the original results, the gray bars show the significant time intervals after adjusting for the subject-wise difference (RAND-REG) in reaction time (mid-gray) and d-prime (light-gray). (**c**) Averaged pupil diameter for all conditions. (**d-f**) Average pupil diameters separated by alphabet size 5, 10 and 15 (left to right), showed sustained larger pupil diameters for random conditions (red, orange and yellow) than regular conditions (shades of blue). (**d**) Alphabet size 5 showed significant differences between REG5 and RAND5 from 2-3s onwards. (**e**) For alphabet size 10, REG10 separates from RAND10 from 3 s onwards with a sustained significant difference from ∼ 7-8 s. (**f**) For alphabet size 15, REG15 separates from REG15 from 4 s, and is significantly different from 6 s onwards. For figures (e) and (f) the significant effects at onset are likely artefacts of regressing out the behavioral measures, resulting from low variability between participants at the onset time points.

#### Pre-processing

Where possible the left eye was analyzed. To measure the pupil dilation response (PDR) associated with tracking the auditory sequence, the pupil data from each trial were epoched from 1 s prior to stimulus onset to stimulus offset (9 s post-onset).

The data were smoothed with a 150 ms Hanning window and intervals where full or partial eye closure was detected (e.g. during blinks) were treated as missing data and recovered using shape-preserving piecewise cubic interpolation. The blink rate was low overall, with the average blink rate (defined as the proportion of excluded samples due to eye closure) at approximately 4% (exp. 1) and 2.6% (exp. 2).

To allow for comparison across trials and subjects, data for each subject in each block were normalized. To do this, the mean and standard deviation across all baseline samples (1 second pre-onset interval) in that block were calculated and used to z-score normalize all data points (all epochs, all conditions) in the block. For each participant, pupil diameter was time-domain averaged across all epochs to produce a single time series per condition.

#### Time-series statistical analysis of pupil diameter

To identify time intervals where a given pair of conditions, REG5 vs RAND5, REG10 vs RAND10, REG15 vs RAND15 exhibited differences in pupil diameter, a non-parametric bootstrap-based statistical analysis was used (Simonoff et al., 1994). Using the average pupil diameter at each time point, the difference time series between the conditions was computed for each participant and these time series were subjected to bootstrap re-sampling (1000 iterations: with replacement). At each time point, differences were deemed significant if the proportion of bootstrap iterations that fell above or below zero was more than 95% (i.e. p < .05). Any significant differences in the pre-onset interval would be attributable to noise, therefore the largest number of consecutive significant samples pre-onset was used as the threshold for the statistical analysis for the entire epoch.

#### Pupil event rate analysis

In addition to pupil diameter, the incidence of pupil dilation events was also analyzed. Pupil dilation events were defined as instantaneous positive sign-changes of the pupil diameter derivative (i.e. the time points where pupil diameter begins to increase).

This activity was analyzed to focus on phasic pupil activity which has been associated with corresponding phasic activity in the Locus Coeruleus and the release of NE (Joshi et al., 2016; Reimer et al., 2016). Following Joshi et al., (2016) and Zhao et al., (2019b) events were defined as local minima (dilations; PD) with the constraint that continuous dilation is maintained for at least 300 ms. For each condition, each subject, and each trial a causal smoothing kernel *ω*(^*τ*^)=*α*^2^×*τ*×*e*^−*αt*^was applied with a decay parameter of *α* = 1/150 ms (Dayan and Abbott, 2001). The mean across trials was computed and baseline corrected. To facilitate the comparison between regular and random sequences, and because pupil dilation events are quite rare (1-2 events per second), we collapsed across alphabet size to derive a single mean time series for REG and RAND. To identify periods in which the event rate significantly differed between conditions, a non-parametric bootstrap-based analysis was used. As for the diameter analysis, this involved computation of a difference time series between conditions for each participant, that was then subject to re-sampling with replacement (1000 iterations). At each time point, differences were deemed significant if the proportion of bootstrap iterations that fell above or below zero was more than 99% (i.e. p < .01).

#### Regressing out behavioral performance

We conducted exploratory analysis to examine whether performance on the incidental gap detection task affected the observed differences in pupil dynamics between REG and RAND patterns. This was achieved by regressing out the variance associated with the gap detection performance from the pupil data. For both experiments each participants’ mean reaction time was used. RT is less limited by ceiling effects and is therefore a good proxy for behavioral difficulty. d prime, was used as a second performance metric for experiment 1. For experiment 2 there were no false alarms and only 5/20 participants were not at ceiling. As a result, it was not appropriate to attempt to model the pupil response to hit rates. To show that these participants were not driving the effect in the pupil we re-ran the main analysis excluding these subjects and present the results in Figure 4-1

**Figure 4:**
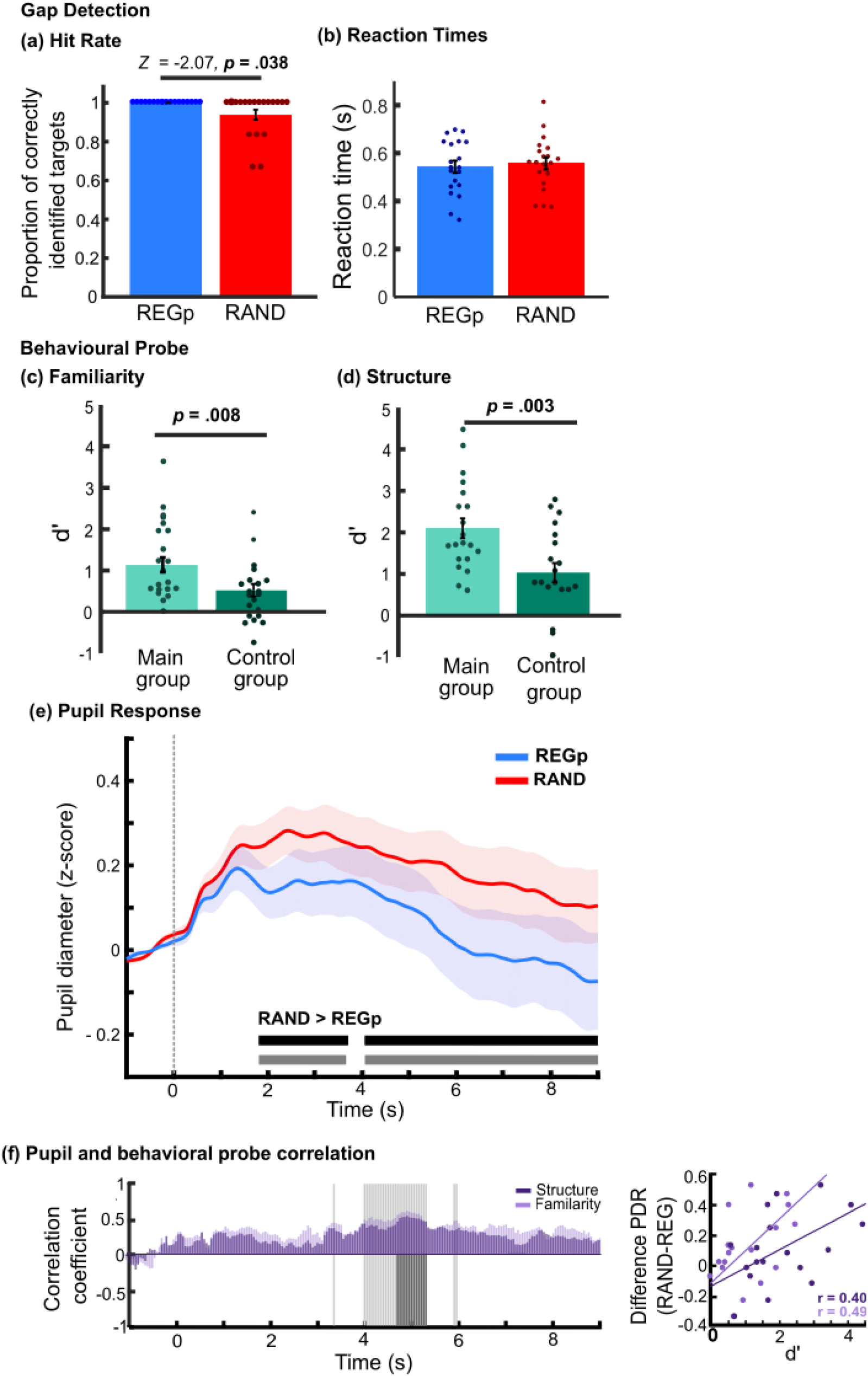
Experiment 2 – probabilistic regularities modulate pupil size and behavior. (a) Hit rate analysis showed more gaps were detected in REGp (blue) than RAND (red) sequences. There were no false alarms (no graph presented). (b) normalized reaction times for gap detection showed no significant differences. Following the main experiment two behavioral probes were separately conducted, in one, listeners were asked to judge if sequences were “familiar (c), and in the other if they contained a “structure (d). D prime (d’) is plotted for the main group (light green) and a control group who had not conducted the main pupillometry experiment (dark green). Error bars show ±1 SEM, circles show individual subjects. (e) Average normalized pupil diameter over time, baseline corrected (−1 – 0s pre-onset). The shaded area shows ±1 SEM. The horizontal bars show time intervals during which significant differences (bootstrap statistics) were observed. The black bar shows the original results, the gray bar shows the significant time intervals after adjusting for the subject-wise difference (RAND-REG) in reaction time. RAND sequences were associated with a larger pupil dilation than REGp from 2 s post-stimulus onset. Also see Figure 4-1, plotting the same data but with the 5 participants with below ceiling performance removed from the analysis (f) Spearman Correlation between the difference in pupil diameter (RAND – REGp) and d’ from the familiarity probe (light purple) and structure probe (dark purple) conducted sample-by-sample (20 Hz) over the entire trial duration. Each purple bar shows the Spearman correlation coefficients at each time point for the two probe tasks. Gray shaded areas indicate time intervals where a significant correlation (p < .05; FWE uncorrected) was observed, light gray corresponds to the correlation with the familiarity probe, significant periods for the structure probe are in dark gray and plotted only on the lower part of the y-axis. For the gray bars, the relationship to the y-axis is for visualization purposes and not meaningful. The plot on the right illustrates the link between pupil size and subsequently assessed sensitivity to regularity by displaying the correlation (Spearman r) between pupil size differences (averaged across 4-6 s) and individual ‘familiarity’ (light purple) and ‘structure’ judgments (dark purple).

Two analysis approaches were taken: the first used average pupil diameter over the latter portion of the trial (4.5 – 9s) where robust differences emerged between conditions (see figures 3d and 4e). Using mean pupil diameter for this time window as the dependent variable, we conducted a repeated measures analysis of covariance (ANCOVA), with a repeating factor of regularity (REG vs RAND) and the difference (RAND-REG) in RT and d’ (experiment 1 only) as covariates. In Experiment 1, this analysis was focused on alphabet size 5 (REG5 vs RAND5), as this showed the most robust effect of regularity on the pupil. To increase power, we also combined the datasets from Experiment 1 and 2, entering Experiment as a between-subjects factor.

The second approach involved regressing out the variance related to the behavioral measures from the unfolding pupil diameter data. For each subject, sample-by-sample differences in pupil diameter (RAND-REG) were regressed onto behavioral performance (difference in RT or d’ between RAND and REG) to remove variance attributable to this potentially confounding factor. The residual pupil data were then analyzed as described in the section Time-series statistical analysis of pupil. This analysis was conducted on all conditions (REG5/RAND5; REG10/RAND10; REG15/RAND15; REGp/RAND in Experiment 2). Because extreme values can skew the regression, the behavioral data were checked for outliers and one participant was removed from the regression analysis with d’ for REG15/RAND15.

## Results

### Experiment 1 – Deterministic regularities

This experiment used sequences of tone pips that were either regularly repeating (REG) or random (RAND; Fig. 1). Previous work showed that brain responses, even from naïve listeners, rapidly distinguished regular from random patterns. The differences emerged as early as 400ms for REG5, 700ms for REG10 and 1050ms for REG15, consistent with the prediction of an ideal observer model which indicated that the emergence of regularity should be detectable from roughly 1 cycle and 4 tones after the introduction of the regular pattern (for details see Barascud et al., 2016; Southwell et al., 2017). Using the same regular sequence structure, we compared the pupil response to regular (REG), highly predictable deterministic sequences to matched random (RAND) sequences of the same alphabet size.

Two factors were manipulated, 1) whether the sequence contained a repeating pattern (REG vs RAND); 2) the alphabet size (5,10 or 15), reflecting the number of different tones in the sequence, and thus its complexity in terms of draw on memory and other perceptual resources.

### Gap detection Results

Sensitivity to the presence of gaps was assessed using d’(Fig 3a). However, as performance was close to ceiling, hit rate and false alarm rate are also presented in Figure 3-1. Parametric tests could not be conducted on d’ due to normality violations, therefore, d’ was initially averaged across alphabet sizes for REG and RAND and compared using a Wilcoxon signed Rank test. This confirmed that d’ was significantly higher for REG (mean = 3.12, std = .50) than RAND (mean = 2.87, std = 0.48, Z = 2.564, *p* = 0.010, Fig 3a). Pairwise Wilcoxon signed rank tests for each alphabet size (Holm-Bonferroni correction was applied) indicated that the effect may be driven by alphabet size 10, as there was a significant difference between REG10 and RAND10 (*Z* = 2.836 *p* = 0.02) but no significant difference between REG5 and RAND5 (*Z* = 1.536, *p* = 0.25) or REG15 vs RAND15 (*Z* = 1.26, *p* = 0.25).

For reaction times (Fig. 3b), a repeated measures (RM) ANOVA with two factors, *Regularity* (REG vs RAND) and *Alphabet size* (5,10,15) revealed a main effect of regularity, with significantly faster response times in REG (mean =0.590 s, SEM = 0.027) compared to RAND (mean = 0.677 s, SEM = 0.031), *F*(1,17) = 41, *p* <.001, ƞp^2^ = 0.71. There was no main effect of alphabet size *F*(2,34) = 0.263, *p* = .771, ƞp^2^ = 0.015, and no interaction *F*(2,34) = 1.786, *p* = 0.183, ƞp^2^ = 0.095.

As an exploratory analysis, we tested whether reaction times varied based on the timing of the gap relative to the sequence onset. As will be demonstrated in the next section, the pupil response to regular sequences emerged later in the trial, particularly for larger alphabet sizes. As we show above, reaction times were faster for REG sequences, therefore we questioned if there were faster reaction times in the latter portion of the trial in the REG condition that were driving both the behavioral effects and pupil response. As each condition only provided 6 target trials, and faster RTs and smaller pupil sizes were observed for all regular conditions, we collapsed across alphabet sizes and calculated the average reaction time for gaps that occurred earlier (< 4.5 s post-sound onset) vs. later in the trial (> 4.5 s post trial onset). An RM-ANOVA was conducted with repeating factor of Time (Early vs Late) and Regularity (REG vs RAND). Reaction times showed a clear effect of regularity (*F* (1,17) = 29.198, p = <.001, ƞp^2^ = .632) but no effect of time (*F* (1,17) = 1.006, p = .316, ƞp^2^ = .059) and no interaction (*F* (1,17) = .009, p = .925, ƞp^2^ = .001).

### Sustained pupil dilation is modulated by sequence predictability

Figure 3c plots the average pupil diameter (relative to the pre-onset baseline) as a function of time. All six conditions share a similar PDR pattern. Immediately after scene onset (t = 0), the pupil diameter rapidly increased, forming an initial peak at ∼0.6 s. Over the next second, pupil diameter slowly increased again to reach a broader peak around ∼3 s after onset. Thereafter, the response entered a sustained phase, which lasted until sequence offset and was associated with a slow continuous decrease in pupil diameter.

Regular sequences elicited a smaller pupil diameter than random sequences, for all alphabet sizes. As can be seen from figure 3, the REG conditions were associated with a faster decrease in pupil diameter (steeper reduction in the sustained response) than the RAND conditions and this effect was modulated by alphabet size. The comparison across matched REG and RAND pairs (Figs. 3d-f) revealed that the separation between traces occurred substantially earlier for alphabet size 5 (Fig. 3d), where a divergence was observed from ∼1.5 s after onset, than the other two conditions. The average trace for REG diverged from RAND at ∼ 3 s for REG10 and ∼4.5 seconds for REG15 (fig. 3e,f) and became statistically significant later in the trial (> 6 s). The staggered divergence is consistent with larger alphabet sizes (i.e. longer REG cycles) requiring more time before a regularity can be established. A similar pattern of divergence latencies has been observed in the brain (Barascud et al., 2016; Southwell et al., 2017), albeit on a faster timescale.

The significant difference between conditions emerged surprisingly late for alphabet size 10, although the conditions separated much earlier. It is likely that a combination of noise and a weaker signal impact the results for this condition.

### Experiment 2 – Non-deterministic patterns

Experiment 2 investigated whether the effects observed in Experiment 1 extend to sequences that contain probabilistic rather than deterministic structure. Towards this aim, we focus on a structure that has been extensively used to study statistical learning in the context of language. Saffran et al., (1996) tested if infants could segment a continuous stream of syllables based only on the statistical regularities between successive items. The streams of syllables had high transitional probabilities within ‘words’ consisting of triplets of syllables, and low transitional probabilities at word boundaries. Infants were found to spend longer looking at non-words that breached the word boundaries, suggesting they had become sensitive to the distributional cues of the syllable stream. Forms of the paradigm have since been used in behavioral and neuroimaging studies (Batterink and Paller, 2017; Farthouat et al., 2017), in adults (Saffran et al., 1997), infants (Saffran, 2020) and other species (Hauser et al., 2001; Toro and Trobalón, 2005) using a variety of stimuli (Saffran et al., 1999; Kirkham et al., 2002). The current experiment uses the pure tone version of this segmentation paradigm (Saffran et al., 1999), with a key modification. The original study used a tone length of 333ms to model the length of syllables, in contrast we use 50ms tones to study this structure at a rate comparable with the sequences in Experiment 1.

To generate the underlying probabilistic structure, twelve different tones were arranged into four tone ‘words’ (see methods). Following Saffran et al. (1999) the same tone ‘words’ were used for each subject. Probabilistic regular sequences (REGp; 9 second-long), generated anew for each trial, were created by randomly ordering the four tone words, with the stipulation that the same tone word could not occur twice in a row (i.e. tone words always transitioned to a different tone word). This created a probabilistic structure where the transitional probability between tones within a word was 1 and the TP at word boundaries was 0.33, see Figure 2 for more details. RAND sequences were generated in the same way as for experiment 1, but using the pool of 12 frequencies from which the tone ‘words’ were created.

The experimental session consisted of three phases. First, participants were familiarized with the REGp sequences. Subsequently, pupil responses were recorded as they listened to REGp or RAND sequences. A gap detection task was used to ensure that participants focused their attention on the sound stream. In a final phase, the same subjects and a control group were asked to make decisions about the familiarity and underlying structure of the different sequence types.

### Gap detection Results

No false alarms were made but there were significantly more gaps detected in REGp compared to the RAND (Wilcoxon Signed Ranks Test: *Z* = 2.07, *p* = .038, Fig. 5a). Reaction times showed no significant difference between conditions (paired samples *t*-test, *t*(19) = -.772, *p* = .450, d = -.173 Fig 4b). Therefore, though the effects are weak and most participants performed at ceiling, the gap detection data demonstrate, similar to Experiment 1, that performance was facilitated in REGp relative to RAND sequences.

**Figure 5:**
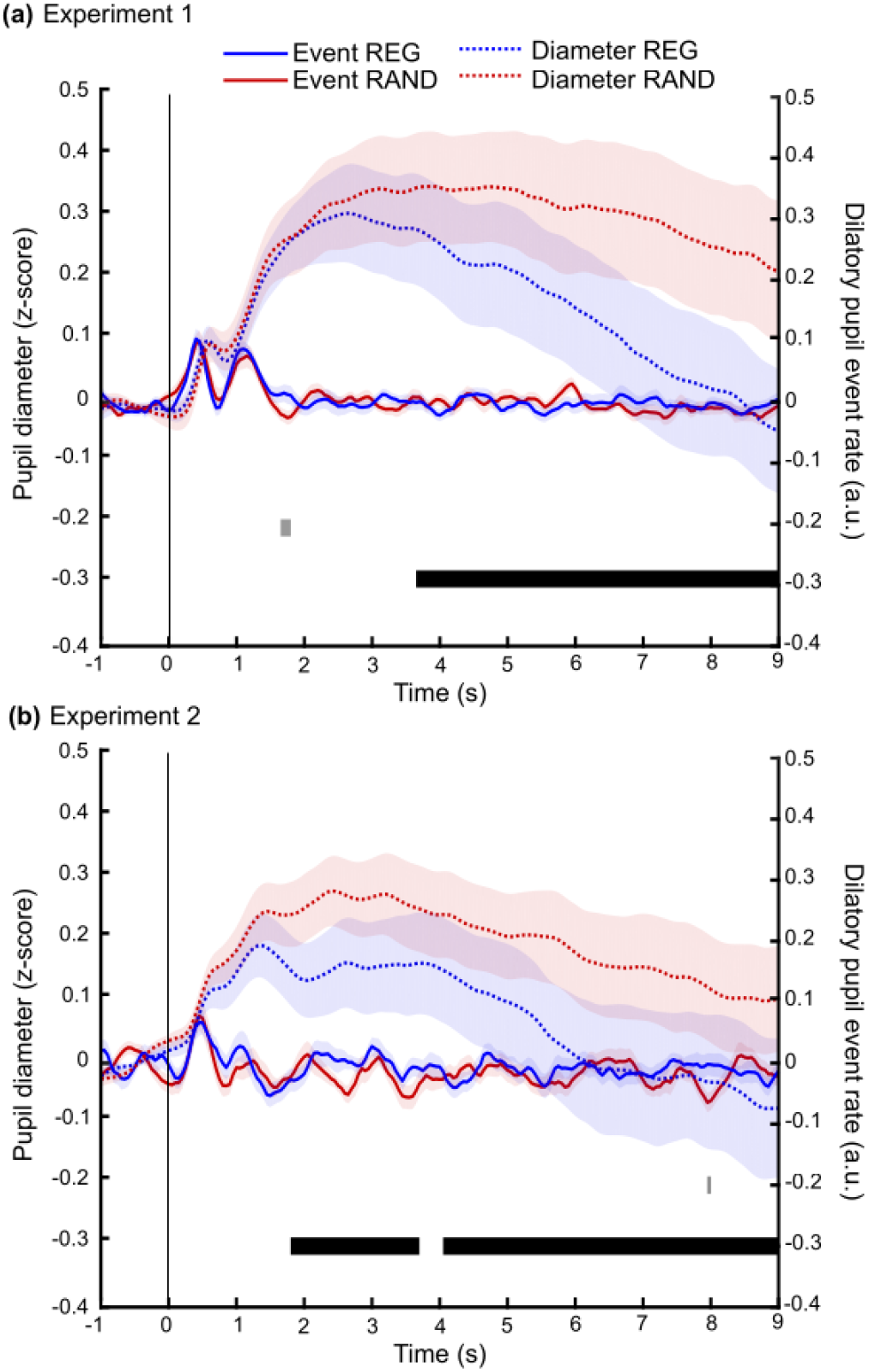
Sequence regularity was not associated with differences in incidence of dilatory pupil events. (**a**) Experiment 1, (**b**) Experiment 2. Solid lines show pupil dilation even rate. Events were defined as the onset of each pupil dilation with a duration of at least 300ms. These were collapsed across alphabet sizes for REG (blue) and RAND (red). Grey markers at the bottom of the graph indicate time intervals where bootstrap statistics showed a significant difference between the two conditions. Dotted lines show the pupil diameter REG (blue) and RAND (red) collapsed across alphabet size. The black bar indicates intervals where bootstrap statistics showed a significant difference between the two conditions. Only the pupil diameter data showed a sustained difference between REG and RAND conditions.

### Exposure to REGp sequences improved subsequent sensitivity to structure

Following the main pupillometry task, participants completed two further tasks, in the first identifying whether a 3-second-long sequence was “familiar” and in the second identifying if the sequence had a “structure” (see methods). These tasks were also completed by a control group who had not participated in the previous phases. The results are shown in figure 4c and d. In both tasks, the majority of participants in the control group showed d’ > 0.This indicates that for some listeners 3 seconds (60 tones) of exposure to the sequence was sufficient to detect a structure, which the listener then interpreted as feeling ‘familiar’. This is in line with previous statistical learning paradigms that show a ‘familiarity’ decision can reflect implicit sequence learning (Forkstam et al., 2008). However, sensitivity in the control group still remained low (d’ < 1) suggesting poor sensitivity overall. Importantly, as expected, the main group showed significantly higher sensitivity than the control group in both tasks (Independent samples t-test, Familiarity: *t*(38) = 2.8, *p* = .008; Structure: *t*(38) = 3.2, *p* = .003), demonstrating that previous exposure improved sensitivity. Unsurprisingly, performance across the ‘familiarity’ and ‘structure’ tasks was correlated for the main (*Spearman’s rho* = .797, *p* < .001) and the control group (*Spearman’s rho* = .570, *p* = .009), confirming that both tasks probed sequence learning (Forkstam et al., 2008).

### Sustained pupil dilation is modulated by sequence predictability

Figure 4e shows the normalized pupil diameter to REG_p_ (blue) and RAND (red) sequences. As in experiment 1, both conditions showed an increase in diameter after sound onset, followed by a sharp decrease in pupil diameter for REGp but not RAND. Since listeners were pre-exposed to the regular stimuli we expected that the pupil response to the REGp condition should rapidly diverge from RAND - as soon as it is statistically possible to differentiate the two sequences (i.e. within 2-3 ‘words’ after sequence onset). Indeed, a sustained difference between conditions emerged from ∼ 2 s post-stimulus onset, roughly at the same time as that observed for REG5 (repeating cycle of 5 tones) in experiment 1. We interpret that as indicating that REGp was differentiated from RAND at a similar latency as REG5 (∼9 tones; see Barascud et al, 2016; Southwell et al, 2017) with the pupil effect manifesting at a delay linked to slower modulatory pathway effects (i.e. the time it takes for the signal to travel from the cortical network which tracks the regularity to the LC and from there to the pupil musculature). However, the extent of divergence between REGp and RAND was smaller than that observed for REG5 (compare 4c and 3d), this was also expected as the probabilistic structure in experiment 2 (see Fig. 2d) retains some degree of unpredictability, i.e. at tone word boundaries. In contrast, REG5 can be predicted with 100% certainty once the tone order has been established.

### Pupil size correlates with (subsequently obtained) explicit identification of structure

An exploratory analysis was conducted into the relationship between pupil dynamics and sensitivity to sequence structure.We correlated the instantaneous PDR difference between REGp and RAND at every time sample (20Hz), with the d’ for each participant (separately for the ‘familiarity’ and ‘structure’ tasks). For this analysis we re-ran the pre-processing to remove blinks without subsequent interpolation to ensure the accuracy of the point by point correlations.

As shown above, performance on the two probe tasks was highly correlated, therefore we expected the two measures to have a similar relationship to pupil diameter. In Figure 4f correlation coefficients (Spearman) are plotted in dark purple (correlation with familiarity probe) and light purple (correlation with structural probe) significant time samples (family-wise error (FWE) uncorrected) are marked in gray, (light gray = familiarity, dark gray = structure). Significant correlations are observed partway through the epoch – between ∼4-6 seconds after onset, revealing that those participants who later indicated high sensitivity to sequence structure were also those exhibiting a larger PDR regularity effect. That correlations appear to be confined to this interval may be due to the fact that the PDR regularity effect stabilizes around that time. The disappearance of correlations towards the end of the trial is consistent with previous observations (Zhao et al., 2019a) and may be because the expectation of trial offset affects pupil dynamics in a manner that interferes with the correlation with behavior.

### Pupil dilation rate is not modulated by predictability

Event rate (instantaneous positive sign-changes of the pupil diameter derivative) was analyzed to focus on phasic pupil activity which has been associated with corresponding phasic activity in the Locus Coeruleus and the release of NE (Joshi et al., 2016; Reimer et al., 2016). To determine whether the observed pupil response is driven by tonic (sustained) or phasic changes in pupil dynamics, we also analyzed the pupil dilation event rate over the course of the trial (see methods). Figure 5 plots both the event rate (solid lines) and dilation response (dotted line) to show how the two measures evolve over time for Experiment 1 (top panel) and Experiment 2 (bottom panel). To improve power in experiment 1, we collapsed across alphabet size, providing a single time series for REG and RAND.

For both experiments the dilation event rate data revealed a series of onset peaks, followed by a return to baseline, with no substantial difference between REG and RAND conditions, in contrast to the robust difference observed for pupil diameter. This suggests that the difference in pupil dynamics between REG and RAND signals is driven largely by tonic rather than phasic pupil activity.

### Regressing out behavioral performance

Both experiments used a gap detection task to ensure that listeners focused their attention on the tone sequence. The task was deliberately easy so as to reduce possible effects of task difficulty on pupil data. However, for some participants regularity was found to modulate performance, increasing the likelihood that a gap would be detected, decreasing the likelihood of making a false alarm (FA) or reducing reaction time (RT). We therefore conducted an additional analysis to confirm that the regularity-linked difference in pupil diameter persists after the variance associated with gap detection performance is regressed out.

Two approaches were taken: According the first, pupil diameter was averaged over the latter portion of the trial (4.5 – 9s) where robust differences emerged between conditions (see figures 3d and 4e). A repeated measures analysis of covariance (ANCOVA) was conducted on pupil size, with a repeating factor of regularity (REG vs RAND) and the difference (RAND - REG) in RT and dprime (d’; experiment 1 only) as covariates. This analysis on experiment 1 data confirmed that the effect of regularity remained significant, F = 7.307, df = 1,15, p = 0.016 ƞp^2^ = 0.328, with no interaction with either covariate, Regularity*RT:*F* (1,15) = 1.635, *p* = .220, ƞp^2^ = 0.098; Regularity vs d’ :*F* (1,15) = .001, *p* = .977, ƞp^2^ = 0. For experiment 2, the ANCOVA could only be conducted with RT as a covariate (see methods). Results confirmed that the effect of regularity persisted: *F* (1,18) = .4.983, *p* = .039, ƞp^2^ = .217 and there was no interaction between regularity and RT:*F* (1,18) = .069, *p* = .796, ƞp^2^ = 004. As a further analysis we also collapsed the data across Experiment 1 (REG5/RAN5) and Experiment 2. As seen above, both yielded similar behavioral effects and pupil dynamics. The ANCOVA confirmed a robust effect of regularity: *F* (1,35) = 15.347, *p* <.001, ƞp^2^ = .968 and no interaction between regularity and RT or experiment (p-values > .2).

A second approach was based on a point-by-point regression analysis. We focused on the subject-wise point-by-point pupil diameter difference between conditions (RAND-REG) and regressed out the behavioral difference between conditions. Statistical analysis (see methods) was then conducted on the resulting time series. The results are plotted (gray horizontal bars) in figures 3d-f and 4e and demonstrate that the main effects of regularity remain after the variance associated with the behavioral variables has been removed.

This experiment was designed to involve a task that ensured the tone sequences were behaviorally relevant. Therefore, there is likely to be a degree of shared variability between performance on the gap detection task and the pupil response to regularity. However, the demonstration that the pupil effects remain after accounting for task performance suggests that effort towards the gap detection task is not driving the pupil effects.

## Discussion

Over two experiments we show that pupil diameter is modulated by the statistical structure of rapidly unfolding auditory stimuli, be they deterministic structures that developed anew on each trial, or more complex statistical structures to which the listener had been pre-exposed. In line with our prediction, we consistently observed a smaller sustained pupil diameter to regular compared with random sequences.

The pupil effects were not correlated with incidental task performance but did reveal a link with subsequently administered familiarity and structure judgements. This demonstrates that pupil dynamics were driven by sequence structure per se, and it’s draw on processing resources, rather than just effort to perform the incidental task.

### Predictability of deterministic sequences modulates sustained pupil size

Previous work has studied pupil responses to deviant stimuli embedded in a predictable structure (Liao et al., 2016; Marois et al., 2018; Quirins et al., 2018; Bianco et al., 2020). Quirins et al., (2018) used a local-global paradigm, also with rapid tone pips. They found that a deviation from the global but not local structure elicited an increase in pupil diameter, but only when actively attending to the deviants, and only in subjects who subsequently showed an awareness of the global regularity. Similarly, Zhao et al., (2019b) showed a transient pupil dilation in response to an unexpected transition from a regular to random pattern. In contrast, the current study examined the dynamics of the pupil response to ongoing regularity.

Participants performed a task that ensured they were broadly attending to the sound sequences. By manipulating the predictability of the tone pip patterns, we were able to assess the extent to which the processing of each sequence type affects pupil-linked arousal.

Based on previous work that demonstrated increased pupil diameter to salient or behaviorally engaging stimuli (Nieuwenhuis et al., 2011; Wang and Munoz, 2015; Liao et al., 2016), we hypothesized that a larger pupil size in response to regular sequences would indicate that attentional resources were engaged to a greater degree by regular relative to random patterns (Zhao et al., 2013). Conversely, a reduction in pupil diameter would indicate that regularity reduces the draw on processing resources by facilitating sequence processing (Southwell et al., 2017). In both reported experiments pupil diameter rapidly decreased once the brain had established the predictable structure of the tone pip sequence, thus supporting the latter hypothesis. In contrast, matched randomly ordered sequences were associated with a largely sustained pupil diameter, suggesting that processing of these stimuli remained more resource-demanding.

For highly predictable, deterministic sequences (Experiment 1), the pupil response showed a rapid divergence between regular and random sequences, reflecting the quick detection of the regular structure. The emergence of regularity was associated with a sustained decrease in pupil size, relative to that evoked by sequences of the same tones presented in a random order. The effect was modulated by alphabet size, with the simplest regular sequences (REG5) showing the more rapid change in pupil diameter.

The pupil response to regularity was consistent with previous neuroimaging work that revealed a rapid change in neural activity following the emergence of regularity (Barascud et al., 2016; Southwell et al., 2017; Herrmann and Johnsrude, 2018).Although the effects seen here arose substantially later than those observed in the brain responses, consistent with a slower pathway (i.e. delays incurred in the pathway between the cortical network that detected the regularity and the pupil). The mechanisms driving the neural response to regularity are poorly understood, but emerging work (Barascud et al., 2016; Auksztulewicz et al., 2017) has implicated an interplay between auditory cortical, inferior frontal and hippocampal sources in the discovery of regularity. A similar network has also been implicated in detecting more complex predictable structure (see Milne et al., 2018 for a summary and also Abla and Okanoya, 2008; Schapiro et al., 2012; Ordin et al., 2020).

### Probabilistic sequence structure modulates pupil size

A clear difference between REG and RAND conditions was also observed for sequences comprised of probabilistic transitions (Saffran et al., 1996). To our knowledge this is the first time that this extensively used structure has been applied to rapid sequences (Saffran et al., 1999). The statistics of the sequence transform it from a stream of individual elements to a series of larger integrated items, in this case triplets of elements, which some argue is a critical component of statistical learning (Batterink and Paller, 2017).

Exploiting this feature of statistical learning, Batterink and Paller (2017) found that as listeners became exposed to the statistical structure they exhibited neural entrainment to not only the rate of individual syllables but also the “words” that were generated using transitional probabilities, (also see Farthouat et al., (2017) for a similar study). Furthermore, there was a correlation between entrainment to the words and reaction time to targets that could be predicted by the structure, supporting a relationship between neural signatures of sequence learning and the influences of sequence learning on subsequent behavior. Critically, we also observed a correlation between modulation of pupil size by sequence type and offline sequence classification (familiarity/structural judgment made after pupillometry measurements), suggesting a relationship between the pupil response to the unfolding sequence and the acquired statistical knowledge; those listeners who showed a larger pupil response difference between REGp and RAND patterns were also those who were better at subsequently discriminating statistically structured from random sequences.

### Predictability modulates tonic rather than phasic pupil activity

Phasic pupil responses (pupil dilation events) have been linked with phasic firing in the LC-NE system (Joshi et al, 2016) and hypothesized to reflect activation of the arousal system. In contrast, slow (tonic) modulation of pupil diameter has been linked to states of perceptual uncertainty (Nassar et al., 2012; Krishnamurthy et al., 2017) and increased demand on processing resources (Sarter et al., 2006). Here, the analysis of pupil dilation event rate demonstrated no difference between conditions, suggesting that the observed pupil effects arise from tonic rather than phasic pupil dynamics.

Krishnamurthy and colleagues (2017) created sequences of sounds played from different locations and asked listeners to make decisions about the locations of upcoming sounds. Over the course of the experiment they manipulated how well the previous sounds could be used to predict the location of an upcoming sound. Where prior information was reliable, the upcoming sound could be accurately predicted. Analysis of baseline pupil dilation, prior to decision making, showed smaller tonic pupil sizes when there were more reliable priors. In other words, as with our data, more predictable stimuli were associated with smaller pupil diameters. Unlike these studies (Nassar et al., 2012; Krishnamurthy et al., 2017), the present results demonstrate sustained changes without perceptual judgements related to the stimulus probability, and with sequences that were too fast for conscious tracking of predictability.

Whilst it may be premature to discuss the underlying brain machinery, the basal forebrain - acetylcholine (BF-ACh) system (Joshi and Gold, 2020) could be hypothesized as a possible underpinning for the observed effects. The basal forebrain has extensive projections in the brain, including to auditory cortex (Guo et al., 2019). Cholinergic signaling has been implicated in the representation of sensory signal volatility (Marshall et al., 2016), and in supporting the rapid learning of environmental contingencies, for example, by boosting bottom-up sensory processing (Yu and Dayan, 2005; Bentley et al., 2011). In the current paradigm the rapid decrease in pupil size during predictable sequences is consistent with a reduction in ACh-driven learning once the sequence structure has been established. A related but mechanistically different proposal is that lower levels of ACh for predictable sequences reflect a decrease in processing demands (Witte et al., 1997; Phillips et al., 2000; Sarter et al., 2006). For REG relative to RAND sequences there is a streamlining of processing that is possible when upcoming tones can be accurately predicted. This contrasts with unpredictable sequences (RAND) where learning cannot take place and thus the resources required to process upcoming tones will remain high.

## Conclusions

We demonstrate that sustained changes in pupil size can be used to identify the emergence of regularity in rapid auditory tone sequences. The results were robust even with a small number of trials (<25 per condition) and consistent across both deterministic and probabilistic sequences. Furthermore, the effects remained after regressing out performance on the incidental task, although future studies may wish to further probe the interactions between the pupil, regularity, and task-related effort. Finally, the speed of sequences used in this paradigm prevented conscious sequence structure tracking, and the task did not require decision making or analysis of the sequence structure. As a result, our findings establish pupillometry as an effective, non-invasive, and fast method to study the automatic extraction of different types of regularities across different populations and even different species.

## Acknowledgements

This work was funded by a Wellcome Trust grant [213686/Z/18/Z] to AEM, a BBSRC grant [BB/P003745/1] to MC and supported by NIHR UCLH BRC Deafness and Hearing Problems Theme.

## Extended Data

**Figure 3-1.**
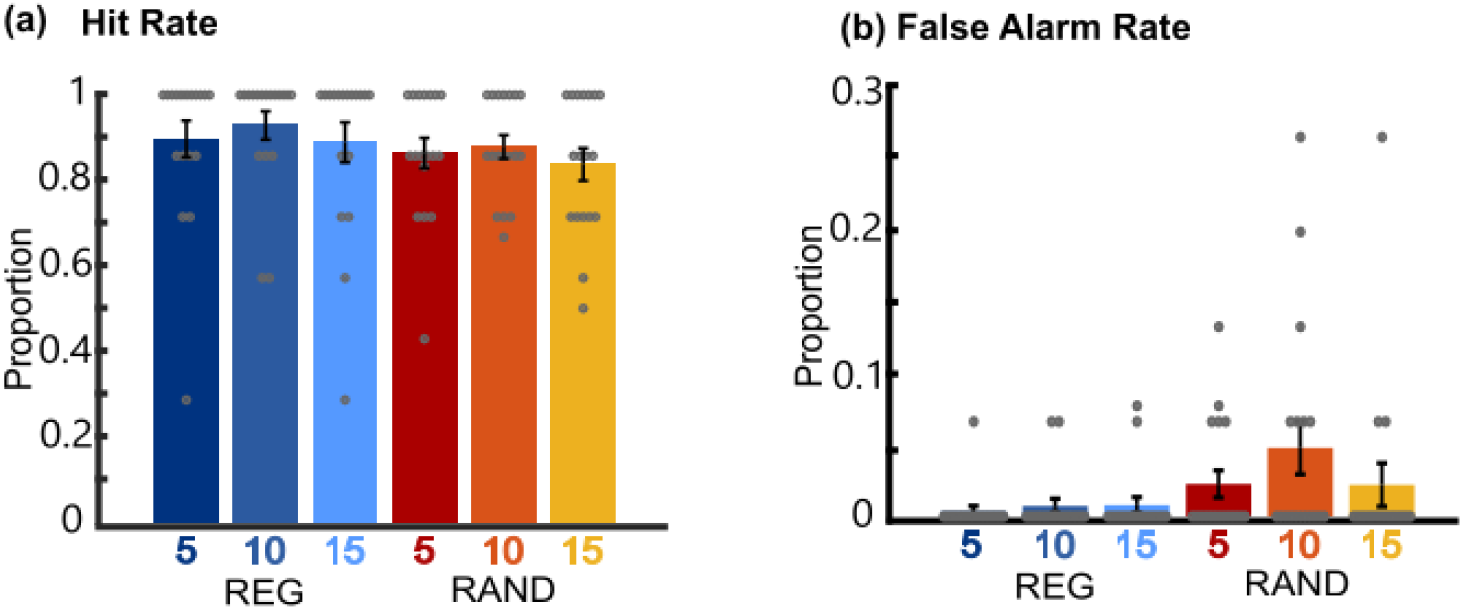
Experiment 1-performance was close to ceiling for the gap detection task. (**a**) hit rate (# hits/ # target trials), (**b**) false alarm rate (# FA / # non-target trials) for experiment 1.

**Figure 4-1.**
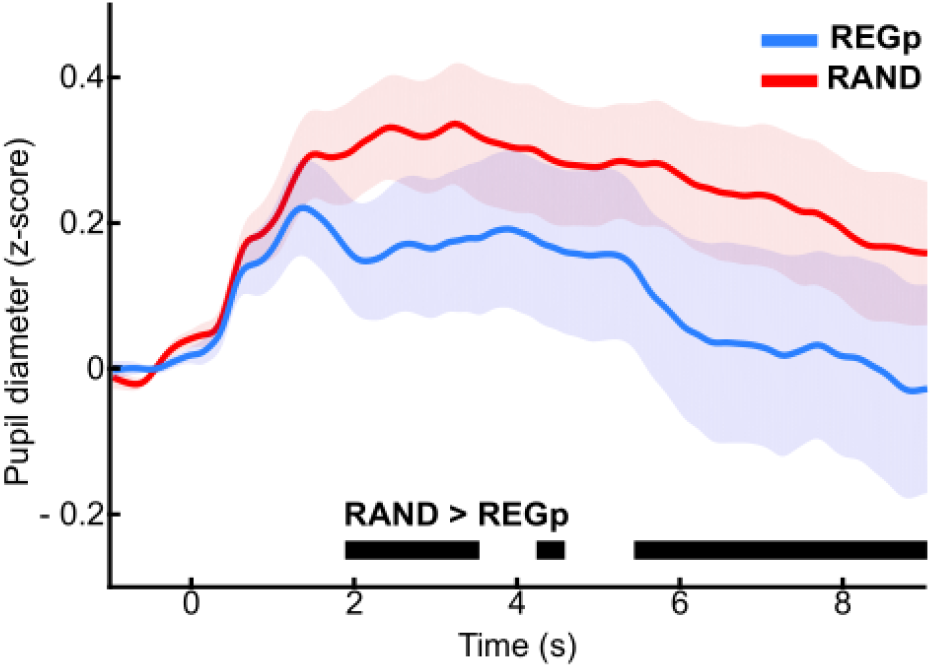
The difference between RAND and REGp remains significant after removal of 5 subjects who exhibited below 100% correct performance in the gap detection task. The figure shows the average normalized pupil diameter over time, baseline corrected (−1 – 0s pre-onset). The shaded area shows ±1 SEM. The black horizontal bar shows time intervals during which significant differences (bootstrap statistics) were observed.

